# Gammaherpesvirus infection triggers the formation of tRNA fragments from premature tRNAs

**DOI:** 10.1101/2024.05.01.592122

**Authors:** Aidan C. Manning, Mahmoud M. Bashir, Ariana R. Jimenez, Heather E. Upton, Kathleen Collins, Todd M. Lowe, Jessica M. Tucker

## Abstract

Transfer RNAs (tRNAs) are fundamental for both cellular and viral gene expression during viral infection. In addition, mounting evidence supports biological function for tRNA cleavage products, including in the control of gene expression during conditions of stress and infection. We previously reported that infection with the model murine gammaherpesvirus, MHV68, leads to enhanced tRNA transcription. However, whether this has any influence on tRNA transcript processing, viral replication, or the host response is not known. Here, we combined two new approaches, sequencing library preparation by Ordered Two Template Relay (OTTR) and tRNA bioinformatic analysis by tRAX, to quantitatively profile full-length tRNAs and tRNA fragment (tRF) identities during MHV68 infection. We find that MHV68 infection triggers both pre-tRNA and mature tRNA cleavage, resulting in the accumulation of specific tRFs. OTTR-tRAX revealed not only host tRNAome changes, but also the expression patterns of virally-encoded tRNAs (virtRNAs) and virtRFs made from the MHV68 genome, including their base modification signatures. Because the transcript ends of several host tRFs matched tRNA splice junctions, we tested and confirmed the role of tRNA splicing factors TSEN2 and CLP1 in MHV68-induced tRF biogenesis. Further, we show that CLP1 kinase, and by extension tRNA splicing, is required for productive MHV68 infection. Our findings provide new insight into how gammaherpesvirus infection both impacts and relies on tRNA transcription and processing.

**Importance:** Diverse conditions of infection and cellular stress incite the cleavage of transfer RNAs, leading to the formation of tRNA fragments which can directly regulate gene expression. In our study of gammaherpesviruses, such as the murine herpesvirus 68 and human oncogenic Kaposi Sarcoma associated Herpesvirus, we discovered that transfer RNA regulation and cleavage is a key component of gene reprogramming during infection. We present the first in-depth profile of tRNA fragment generation in response to DNA virus infection, using state-of-the-art sequencing techniques that overcome several challenges with tRNA sequencing. We present several lines of evidence that tRNA fragments are made from newly-transcribed premature tRNAs and propose that this may be a defining characteristic of tRNA cleavage in some contexts. Finally, we show that tRNA splicing machinery is involved with the formation of some MHV68-induced tRNA fragments, with a key regulator of splicing, CLP1, required for maximal viral titer. Together, we posit that tRNA processing may be integral to the elegant shift in gene expression that occurs during viral take-over of the host cell.

## Introduction

Transfer RNAs (tRNAs) are important non-coding RNAs with dual functionality, essential for protein translation and, when processed, as rich sources of functional small RNAs in the overall pool of tRNA fragments (tRFs). Advances in small RNA sequencing technology have revealed an abundance of tRFs made by endonucleolytic cleavage of tRNAs, which typically increase in abundance in response to cellular stress, viral infection, and other disease states^1–6^. While some cleavage products may solely be degradation intermediates, some tRFs have been shown to have protein binding partners that would impart functionality, such as Argonaute or Ybx1^7, 8^. Diverse classes of tRFs can be made from tRNAs—fragments of differing lengths can be generated from the 5’ end, 3’ end, or internal region of the tRNA, with sequence diversity arising from the parental tRNA family origin. The diversity of sequence and structure has resulted in overlapping naming systems for tRFs, including tRNA halves, tRNA-derived RNAs (tDRs)^9^, tiRNAs^10^, among others. Notably, tRF diversity explains the numerous functions and protein interactors observed for tRFs^7, 8, 11^. Comprehensively identifying tRFs produced in response to stress and infection is fundamental to understanding the role tRFs play in different cellular responses.

Viral infection involves an overhaul of host gene expression and thus represents a system in which tRNA regulation likely plays a significant role. We have previously explored the tRNA expression landscape during infection using murine gammaherpesvirus 68 (MHV68), which is a large double-stranded DNA virus genetically similar to two cancer-causing human viruses, Kaposi sarcoma associated herpesvirus (KSHV) and Epstein-Barr virus (EBV)^6^. Due to strict human tropism of KSHV and EBV, MHV68 has served as an important model to understand gammaherpesvirus pathogenesis and host antiviral control^12^. MHV68 uniquely encodes 8 viral tRNA-miRNA encoded RNAs (TMERs) that are important for pathogenesis^13, 14^. Each TMER encodes a viral tRNA (virtRNA) followed by one or two pre-miRNA stem-loops. Expression of the viral miRNAs require host tRNA transcription and processing machinery, including RNA polymerase III and tRNAseZ/ELAC2^13–15^. Because of its genetic similarity to human gammaherpesviruses and its exploitation of tRNA expression machinery, we reasoned that using MHV68 infection serves as an ideal model for elucidating novel mechanisms of tRNA transcription and processing in mammalian cells. Our previous tRNA profiling of MHV68-infected fibroblasts demonstrated selective upregulation of approximately 20% of host tRNA genes; however, no patterns regarding specific tRNA families emerged^6^. Additionally, differential expression of tRNAs during infection manifested primarily at the level of premature tRNA (pre-tRNA) transcripts with minor changes in mature tRNA pools (<2-fold). This work implied that the efficiency of tRNA processing and maturation is decreased during MHV68 infection, but technical limitations of the sequencing method employed (DM-tRNA-seq^6, 16^) hindered our ability to explore shorter derivatives of tRNAs in the sample.

Mounting evidence suggests that tRFs can functionally impact the replication of diverse RNA viruses, including retroviruses, respiratory syncytial virus, and hepatitis C virus, through binding and regulating viral and host RNAs and proteins^2, 4, 17^. Inspired by these emerging indications of tRF roles in virally infected cells, we profiled host and viral tRFs produced in response to DNA virus infection. We applied recently described library preparation and tRNA mapping methodologies, in a strategy we refer to here as OTTR-tRAX, to small RNA isolated from MHV68-infected murine fibroblasts. OTTR-tRAX recapitulated the pre-tRNA upregulation in response to MHV68 infection^6^. It also revealed widespread tRF production from both for host and viral tRNAs (virtRNAs) as a molecular signature of gammaherpesvirus infection. Surprisingly, we found that many tRFs are derived from pre-tRNAs. Some pre-tRFs match expectation for tRNA splicing intermediates. Indeed, we link their presence to the tRNA splicing endonuclease, TSEN2, as well as the tRNA splicing regulatory RNA kinase, CLP1. We show that TSEN2 and CLP1 promote the generation of 5’ pre-tRFs derived from pre-tRNA-Tyr. Further, we show that CLP1 is required for efficient MHV68 replication. Altogether, we provide evidence that tRNA regulation by processing impacts DNA virus infectivity.

## Materials and Methods

### Cell culturing conditions

NIH 3T3 murine fibroblasts, NIH 3T12 murine fibroblasts, and MC57G fibrosarcoma cells (ATCC) were maintained in high-glucose Dulbecco’s modified Eagle’s medium (DMEM) supplemented with 10% fetal bovine serum (FBS). Cultures were screened regularly for mycoplasma (Invivogen) and cultured in the absence of antibiotics.

### Virus infections

Green fluorescent protein (GFP) expressing MHV68-R443I carrying a single point mutation in the viral endonuclease muSOX and the MHV68-MR mutant revertant were amplified in NIH 3T12s^18, 19^. The MHV68-MR was used as the source of wild-type MHV68 for all experiments in this work. MHV68-infected NIH 3T12 cells and media were subjected to two freeze-thaws, then cellular material was pelleted and discarded. Viruses present in the supernatant were pelleted at high-speed (12K x g) for 2 h, then treated with DNase (Fisher #EN0523) for 1 h at 37 °C. Media was added and virus was pelleted at high-speed (12K x g) for 2 h. Virus pellet was dissolved in DMEM +10% FBS media, aliquoted, and stored at −80 °C. MHV68-GFP titer was measured on NIH 3T3 cells for the 50% tissue culture infective dose (TCID50). For high MOI/single-step infection, MHV68 virus was added to ½ culture volume of media, placed on NIH 3T3 or MC57G cells as indicated for 1 h at multiplicity of infection (MOI) of 5 to allow viral entry, then replaced with fresh DMEM +10% FBS medium.

### siRNA treatment

Pooled siRNA purchased from Dharmacon was nucleofected into 5×10^5^ NIH 3T3 cells at 200nM. The Neon™ Transfection System (Thermo Fisher), with buffer E2 and 100uL tips, was used to perform the nucleofection (1400 volts/20 ms/2 pulses). Cells were infected 24 h post-nucleofection and harvested 24 h post-infection (hpi). Total RNA was analyzed via RT-qPCR.

### RT-qPCR and SL-qPCR

Total RNA was isolated from cells using TRIzol (Invitrogen) and was treated with Turbo DNase (Ambion). Turbo-treated RNA was then reverse transcribed with AMV RT (Promega) primed with random 9-mer (IDT) for total RNA measurements by RT-qPCR or stem-loop (SL) primers for 5’ tRF measurements by SL-qPCR. Binding of SL primers requires an exact 3’ end for successful amplification, thus we designed primers using the most frequent 3’ end of upregulated 5’ tRFs in our dataset. Our SL-qPCR assays will detect the following using tRFnamer^9^ nomenclature using mm10 as a sequence source: tDR-1:37-Tyr-GTA-1-M2, tDR-1:17-Asn-GTT-1, and tDR-1:24-Gln-CTG-1-M3. Gene expression was measured by amplifying cDNA using iTaq SYBR Green Supermix (Biorad) on a QuantStudio 3 (Applied Biosystems), and all primers are listed in Table 1. RT-qPCR was analyzed using the delta delta CT (ΔΔCT) method. Average fold changes were calculated relative to 18S or U6 internal controls.

**Table 1.**
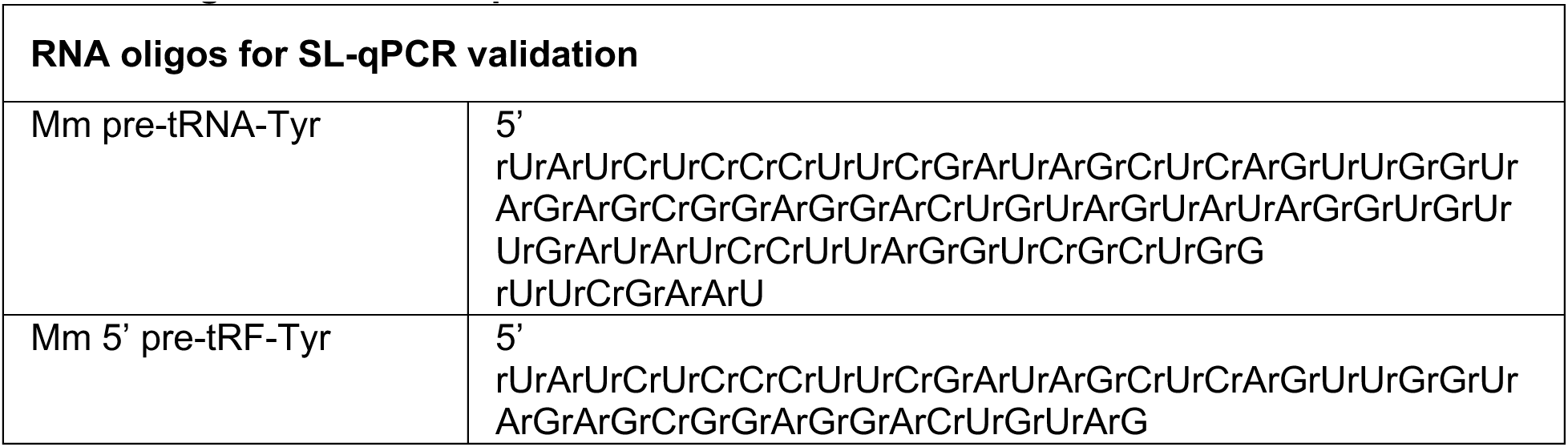
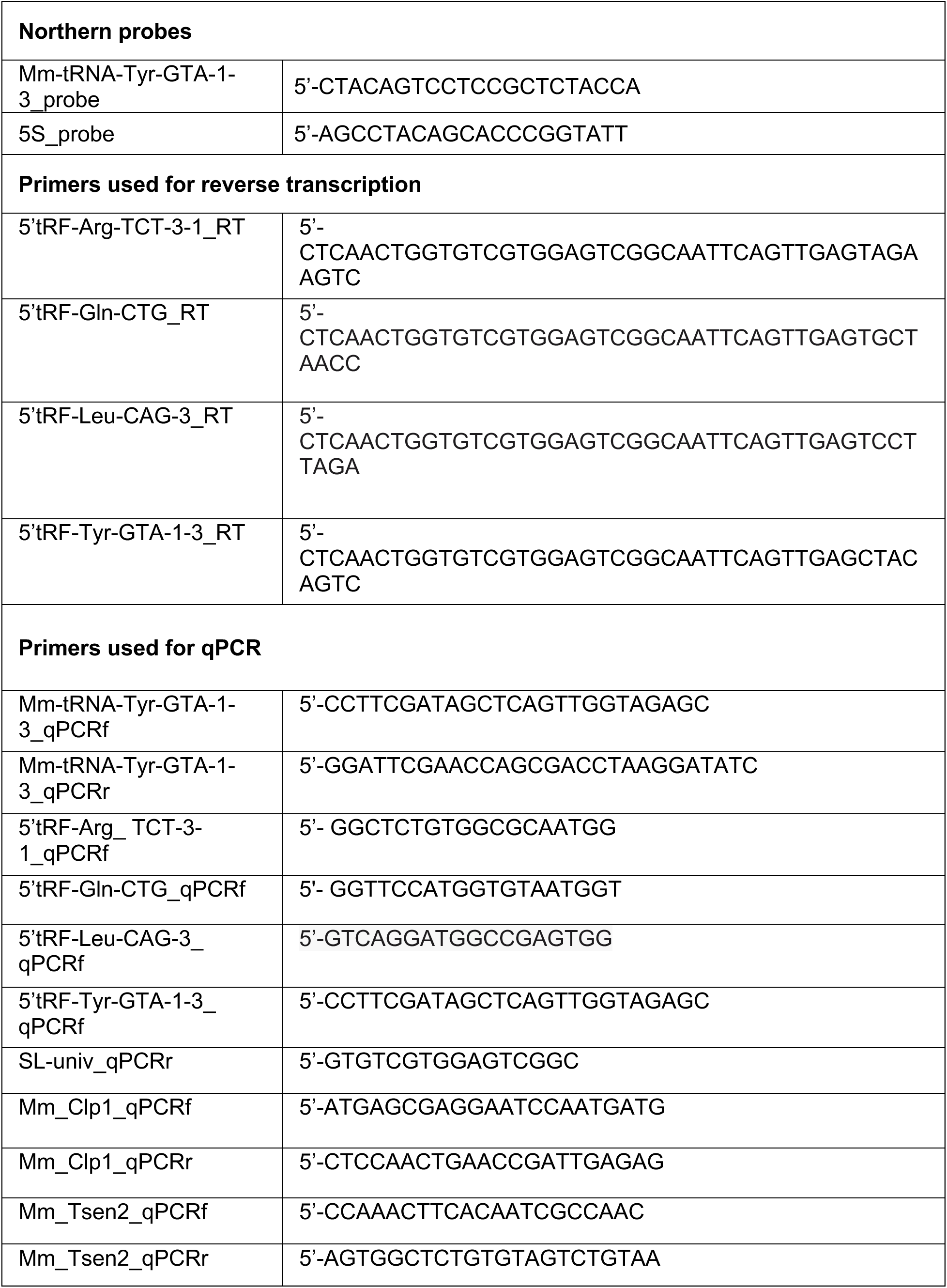
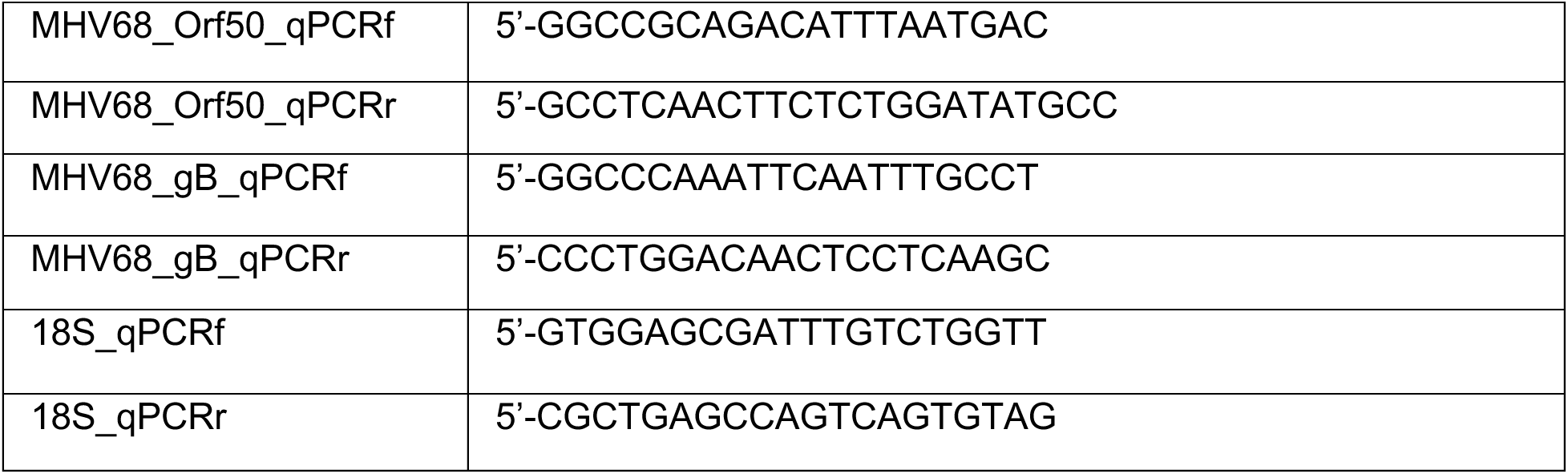
Oligonucleotide sequences.

### Northern analysis

Total RNA was loaded onto 8 to 12% PAGE–7 M urea gels and transferred to Hybond-N+ (GE) membranes using a Trans-Blot Turbo Transfer system (Bio-Rad) in 1X TBE. Blots were crosslinked using the auto-crosslink setting on a UV Crosslinker FB-UV-XL-1000 (1,200 μJ × 100) (Fisher Scientific) and prehybridized in ULTRAhyb buffer (Thermo Fisher) at 42°C for 1hr before adding radiolabeled probe. Probes were generated by end labeling oligonucleotides specific for Tyr or 5S (Table 2) as a loading control using T4 PNK and [γ-32P] ATP. Blots were probed overnight at 55°C and washed 3 times for 10 minutes in 0.5× SSC (1× SSC is 0.15 M NaCl plus 0.015 M sodium citrate) and 0.1% SDS. Blots were imaged using a Typhoon imager and processed on Photoshop (Adobe). Densitometry was performed using BioRad ImageLab software.

### Ordered two template relay sequencing (OTTR-seq)

Small RNAs were isolated from mock-infected, MHV68-MR-infected, or MHV68-R443I-infected MC57G cells (MOI=5, 24 h) in biological triplicate using miRVANA isolation kits (Thermo Fisher) and was used for generating OTTR-seq libraries^20^. Briefly, deacylated, CIP-treated, and PNK-treated small RNA was 3’ tailed using mutant BoMoC RT in buffer containing only ddATP for 90 minutes at 30 °C, with the addition of ddGTP for another 30 minutes at 30 °C. This was then heat-inactivated at 65 °C for 5 minutes, and unincorporated ddATP/ddGTP were hydrolyzed by incubation in 5 mM MgCl2 and 0.5 units of shrimp alkaline phosphatase (rSAP) at 37 °C for 15 minutes. 5 mM EGTA was added and incubated at 65 °C for 5 minutes to stop this reaction. Reverse transcription was then performed at 37 °C for 30 minutes, followed by heat inactivation at 70 °C for 5 minutes. The remaining RNA and RNA/DNA hybrids were then degraded using 1 unit of RNase A at 50 °C for 10 minutes. cDNA was then cleaned up using a MinElute Reaction CleanUp Kit (Qiagen). To reduce adaptor dimers, cDNA was run on a 9% UREA page gel, and the size range of interest was cut out and eluted into gel extraction buffer (300mM NaCl, 10mM Tris; pH 8.0, 1mM EDTA, 0.25% SDS) and concentrated using EtOH precipitation. Size-selected cDNA was then PCR amplified for 12 cycles using Q5 High-fidelity polymerase (NEB #M0491S). Amplified libraries were then run on a 6% TBE gel, and the size range of interest was extracted to reduce adaptor dimers further. Gel slices were eluted into gel extraction buffer (300mM NaCl, 10mM Tris; pH 8.0, 1mM EDTA) followed by concentration using EtOH precipitation. Final libraries were pooled and sequenced using 150 SE and 200 cycle kit on an Illumina NovaSeq.

### tRNA Analysis of eXpression (tRAX)

Sequencing adaptors were trimmed from raw reads using cutadapt, v1.18, and read counts were generated for host and viral small RNA types using tRAX^21^. Briefly, trimmed reads were mapped to a combined mouse (GRCm38/mm10) and gammaherpesvirus (MHV68) reference containing mature tRNAs obtained from GtRNAdb^22^ and their corresponding genomic sequences (for mapping sequences derived from pre-tRNAs) using Bowtie2 in very-sensitive mode with the following parameters to allow for a maximum of 100 alignments per read: --very-sensitive --ignore-quals --np 5 -k 100. Mapped reads were filtered to retain only the “best mapping” alignments. Raw read counts of tRNAs and other small RNA types were computed using tRNA annotations from GtRNAdb, and annotations from GENCODE M23 and miRBase v22^23^. A full description of the mapping parameters for tRFs can be found here^21^. In brief, reads are first mapped to a custom-built reference database for mature tRNAs, including the fully-processed tRNA sequences with 3’ CCA tails. For mature tRNA reads, if the read covers both the 5’ and 3’ boundaries of the mature tRNA, it is considered a full-length mature tRNA. For tRFs, tRAX computes separate read counts for tRFs that are within 10 nt of the 5’ end, 3’ end, or internal sequences that are not within 10 nt of either end (“other’) of mature tRNA sequences. In contrast, if sequencing reads have at least 5 nt that extend beyond the 5’ and 3’ boundaries of the mature tRNA (only one extension is required), they map best to the genomic database and are binned as pre-tRNAs. Within pre-tRNA reads, if both transcript ends extend at least 2 nt beyond the 5’ and 3’ boundaries, the read is assigned as a full-length pre-tRNA. Otherwise, it is assigned as a partial pre-tRNA. Partial pre-tRNAs are not further binned, so the 5’ pre-tRFs discussed in this manuscript were manually extracted and identified using tDRnamer^9^. We note that it is not possible to distinguish whether tRFs binned as “5’”, “3’”, or “other” came from a premature or mature tRNA, unless an intron or 3’ CCA sequences are present. Raw read counts were then normalized and compared using DESeq2 and its associated statistical tests. Data has been deposited with NCBI GEO, identification number: GSE255627.

### Statistical analysis

All experiments were performed in at least biological triplicate, meaning that experiments were performed on different days with different cell populations in terms of stock vial or passage number. All RT-qPCR statistical analysis was performed using raw ΔCt values. The statistical test used is indicated in the figure legends.

### Reagents

**Reagents:**
|  | **Description** | **Catalog Number** |
| --- | --- | --- |
| **Cell culture** | DMEM, high glucose | Gibco 11965092 |
|  | DPBS, no Ca/Mg | Gibco 14190250 |
|  | Fetal Bovine Serum | Gibco 26140079 |
|  | Trypsin-EDTA (0.25%) | Gibco 25200056 |
| **RT-qPCR** | SuperScript III RT | Thermo Fisher 18080044 |
|  | AMV RT | Promega M5108 |
|  | MgCl<sub>2</sub> | Bio-Rad 1708872 |
|  | Turbo DNase | Invitrogen AM2239 |
|  | 10mM dNTPs | Thermo Scientific FERR0193 |
|  | RNAasin | Promega N2515 |
|  | iTaq Universal SYBR Green Supermix | Bio-Rad 1725125 |
| **Instruments** | Neon transfection system | Thermo Fisher MPK5000 |
|  | QuantStudio 3 | Applied Biosystems A28131 |
|  | Proflex thermocycler | Applied Biosystems 4483636 |

### Biological Resources

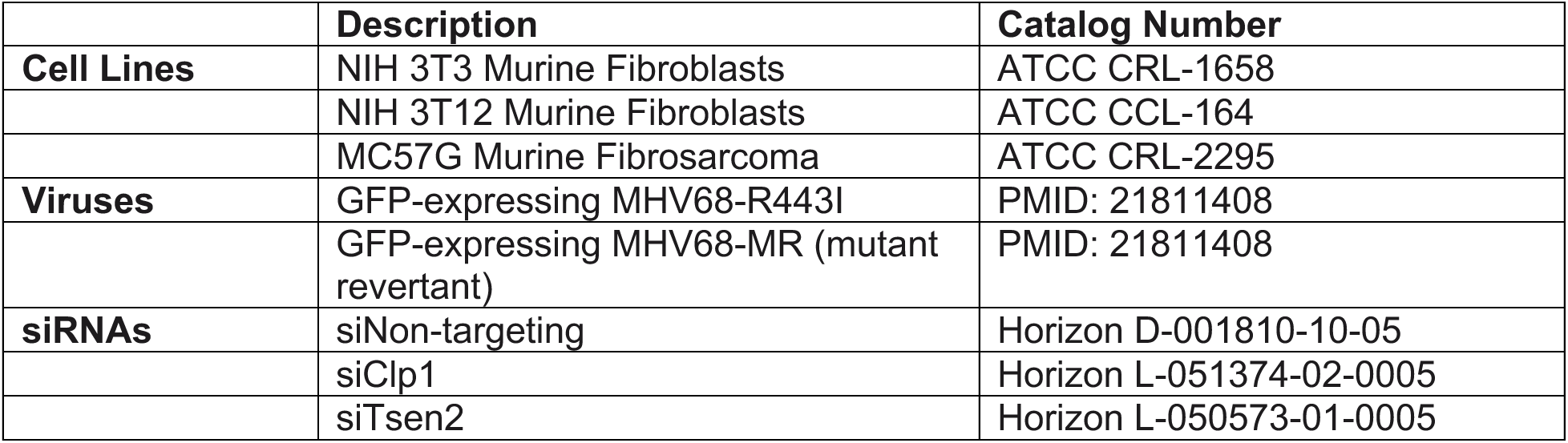

## Results

### Quantitative profiling of tRNAs and tRFs during MHV68 infection

To quantitatively profile tRNA and tRF expression during MHV68 infection, we considered new methodologies to enhance the accuracy and throughput of combined tRNA/tRF sequencing, which is inherently difficult due to both technical and bioinformatic challenges. Standard small RNA library preparations use retroviral reverse transcriptases (RTs) that lack sufficient processivity to reverse transcribe highly modified and structured tRNAs and tRFs from end to end (Fig 1).^24^ Coverage of full-length tRNAs is dramatically improved with the use of more processive RTs, such as TGIRT^25^ used in DM-tRNA-seq^16^. TGIRT uses a template-jumping mechanism to initiate reverse transcription, adding the first sequencing adaptor during cDNA production. This eliminates the need to ligate adaptors prior to reverse transcription, which is convenient for low RNA input and workflow. However, this workflow allows partial cDNA products resulting from disengagement of the RT at modified bases into the sequencing pool; thus, the incomplete cDNA products are indistinguishable from biologically-relevant tRFs. Because of these technical issues, it is current standard practice to analyze full-length tRNAs and tRFs using separate library preparation techniques, such as DM-tRNA-seq for full-length tRNAs and ARM-seq for tRFs^16, 26^. To overcome these challenges and facilitate a single approach to monitor both tRNAs and tRFs by sequencing, we applied Ordered Two Template Relay (OTTR)^20^ to prepare tRNA sequencing libraries. OTTR harnesses the template jumping activity and high processivity of an engineered eukaryotic retrotransposon RT, called BoMoC, to sequentially add both adapters to RNA (rather than only one adaptor with TGIRT)^20, 27^. Processivity through the length of the tRNA or tRF is required for the 2^nd^ adaptor addition (Fig. 1). Thus, this technology is well-suited for simultaneous tRNA and tRF analysis, as it should minimize *and* exclude partial cDNA synthesis products from the sequenced pool. We have combined this technology with best-practice and user-friendly tRNA and tRF mapping software, called tRAX^21, 28^, into a pipeline we refer to here as “OTTR-tRAX”.

**Figure 1.**
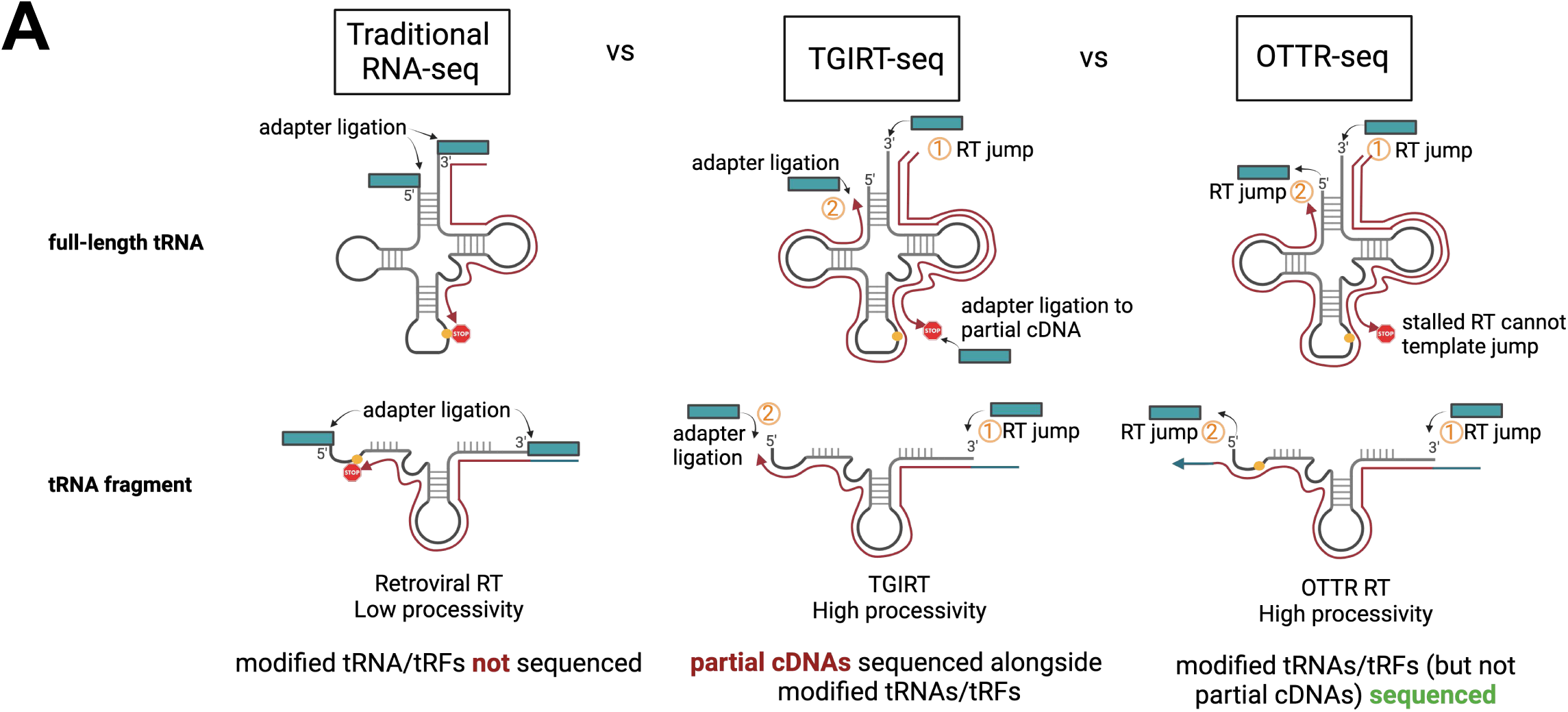
Benefits of OTTR-seq for tRNA sequencing. Traditional RNA-seq uses retroviral reverse transcriptases (RTs) with low processivity through tRNA modifications, resulting in depletion of modified transcripts in final sequencing libraries. TGIRT-seq uses the more processive TGIRT enzyme, but allows partial/incomplete cDNA products into library assembly where they are indistinguishable from biologically-relevant tRFs. The high processivity and template jumping activity of the OTTR RT used in OTTR-seq ensures that highly modified tRNAs and genuine tRFs (and not partial cDNA products) are sequenced.

We performed OTTR-tRAX on small RNA extracted in biological triplicate from the murine fibroblast cell line MC57G with or without a single round of infection with MHV68 (24 hours) at an MOI=5, conditions we have previously analyzed by DM-tRNA-seq^6^. We performed steps to deacylate (treatment at pH=9), remove 5’-P (CIP treatment), and remove 3’ cyclic-phosphates (PNK treatment) from tRNAs and tRFs to offset potential interference of these modifications during cDNA library preparation. To account for the known bias of PCR to amplify shorter fragments, the final cDNA library was size selected, PCR amplified, and sequenced in separate pools corresponding to 15-50 nt or 50-200 nt inserts. We analyzed our sequencing data using the tRAX software package for the analysis of tRNAs and tRFs (see Methods)^21^. More than 60% of the OTTR reads in the 50-200 nt size class were from tRNAs (green) (Fig 2A, right top 2 bars). In comparison, our previous DM-tRNA-seq library contained three-fold less reads mapping to tRNAs (20%) and was instead enriched for snoRNAs (tan) at ∼50% of mapped reads^6^ (Fig 2A, right bottom 2 bars). The composition of the 15-50 nt size class library using OTTR was derived from a variety of sources, including tRFs (green), which comprised the largest percentage at 30-40%, but also included sequences mapping to rRNAs (orange), miRNAs (pink), and snoRNAs (tan). We also assessed how well OTTR-tRAX performed on full length mature tRNAs. To assess the levels of full-length mature tRNAs represented in our OTTR libraries, we compared normalized read counts of each tRNA isodecoder using a read-length cutoff of greater than 70 nucleotides to analyze full-length tRNA transcripts (Fig 2B). Most isodecoders had at least one representative with expression over the median read count of all measured tRNAs (horizontal dotted line), indicating a diverse pool of tRNA species were captured. Importantly, OTTR-tRAX was robust enough to recapitulate the upregulation of viral and host pre-tRNA species during MHV68 infection we reported prior (Fig 2C)^6^.

**Figure 2.**
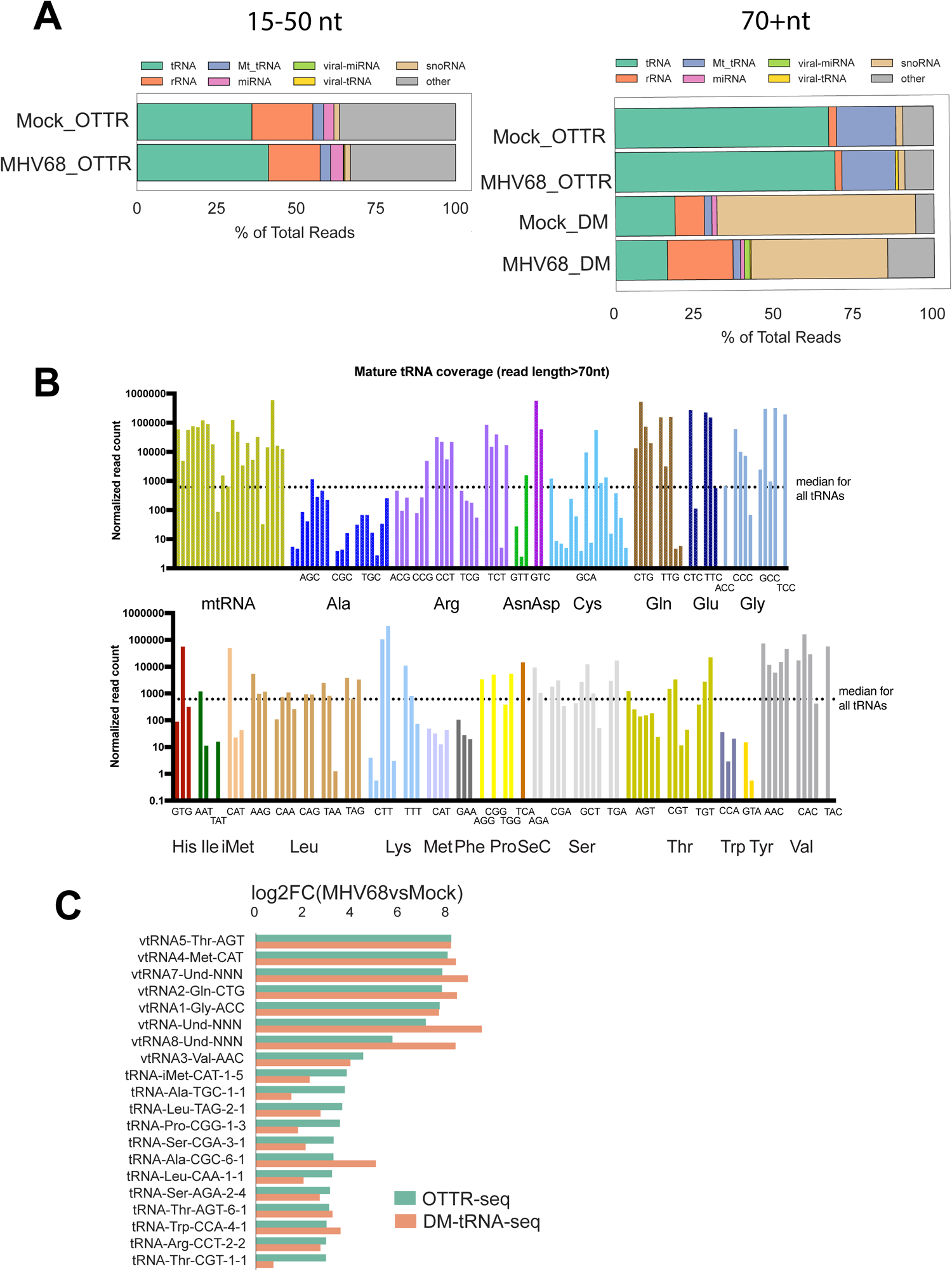
OTTR-seq robustly captures host tRNAs. (A) Distribution of reads mapping to various small RNA classes using OTTR-seq (OTTR) or DM-tRNA-seq (DM) prepared from mock- and MHV68-infected MC57G mouse fibroblasts at an MOI=5 for 24 hours. (B) Normalized read counts for host tRNA isodecoders detected in mock-infected libraries using OTTR-seq. The dotted line marks the median normalized read count for all tRNA isodecoders as reference. (C) Log2(fold-change) values between MHV68-infected (MR) versus mock infection from OTTR-seq (green) or DM-tRNA-seq (orange) are consistent for viral-tRNAs (“virtRNA” prefix) and host pre-tRNAs.

### MHV68 infection causes host shutoff-dependent changes in tRNA expression

MHV68-induced “host shutoff,” in which cellular mRNA is endonucleolytically cleaved by the viral endonuclease muSOX (ORF37), contributes to the accumulation of pre-tRNAs in infected cells^6^. Importantly, muSOX does not cleave reporter transcripts made by RNA polymerase I or II^29^, arguing against the possibility of cleavage of tRNAs by muSOX. This is further supported by the fact that MHV68-R433I, containing a single point mutation in muSOX with diminished mRNA endonucleolytic activity,^19^ lowers the levels of host pre-tRNA-Tyr and - Leu compared to mutant revertant, MHV68-MR, as measured by RT-qPCR^6^. We previously hypothesized that global mRNA degradation during host shutoff depletes the cell of post-transcriptional tRNA processing factors required for pre-tRNA maturation and turnover, extending the half-life of pre-tRNAs^6, 30^. Together, this data suggest that muSOX activity contributes to pre-tRNA accumulation during infection.

To more extensively profile pre-tRNAs differentially affected during infection with MHV68-R443I, MC57G mouse fibroblasts were mock-infected or infected with MHV68-R443I or MHV68-MR virus at an MOI=5 for 24h in biological triplicate. Notably, the MHV68-R443I virus has identical replication kinetics to MHV68-MR in cultured fibroblasts^19^. We first confirmed similar levels of infection with MHV68-R443I or MHV68-MR by measuring the level of viral ORF50 transcript expressed at 24 hours post infection (Fig 3A). Host shut-off activity was assessed by measuring the level of the host transcript, *Gapdh*. As expected, MHV68-MR infection leads to decreased *Gapdh* expression, while MHV68-R443I restored expression to that observed in mock-infected cells (Fig 3A). Mock-infected, MHV68-MR-infected, and MHV68-R443I RNA samples were then processed using OTTR-tRAX (Fig 3B). Differentially expressed (p<0.05) pre- or mature tRNAs from the 50-200 nt size class upon MHV68-MR infection compared to mock are depicted in Fig 3B (MHV68-MR vs. Mock). The majority of significantly (p<0.05) differentially expressed transcripts are premature tRNA sequences, as seen previously using DM-tRNA-seq^6^. MHV68-R443I infection also increased pre-tRNA expression, but to a lesser degree, which might be due to either residual host shutoff activity or additional contributing factors (Fig 3B, MHV68-R443I vs. Mock). However, the majority of pre-tRNAs exhibited decreased expression in MHV68-R443I compared to MHV68-MR (Fig 3B, MHV68-R443I vs. MHV68-MR), confirming that pre-tRNA accumulation is enhanced by muSOX activity^6^. Read coverage plots of pre-tRNA-Tyr-GTA-1-3 and pre-tRNA-Gln-CTG-1-1 illustrate the reduced pre-tRNA expression with MHV68-R443I versus MHV68-MR virus (Fig 3C). There were no differences in the lengths of the 5’ leaders or 3’ trailers of pre-tRNAs between mock- and MHV68-infected cells, suggesting that these transcripts correspond to nascent pre-tRNAs prior to trimming or splicing. Differential upregulation is specific to a subset of pre-tRNAs (Fig 3B), as other non-coding RNAs made by RNA polymerase III do not increase in abundance during infection with MHV68-MR or -R443I (Fig 3D). We posit that elevated host tRNA/tRF levels in infected cells is an indirect effect of multiple downregulated host pathways, starting with the depletion of tRNA maturation or turnover machinery. Overall, OTTR-tRAX reveals global changes in pre-tRNA abundance that are partially dependent on the host shutoff functionality of MHV68 muSOX.

**Figure 3.**
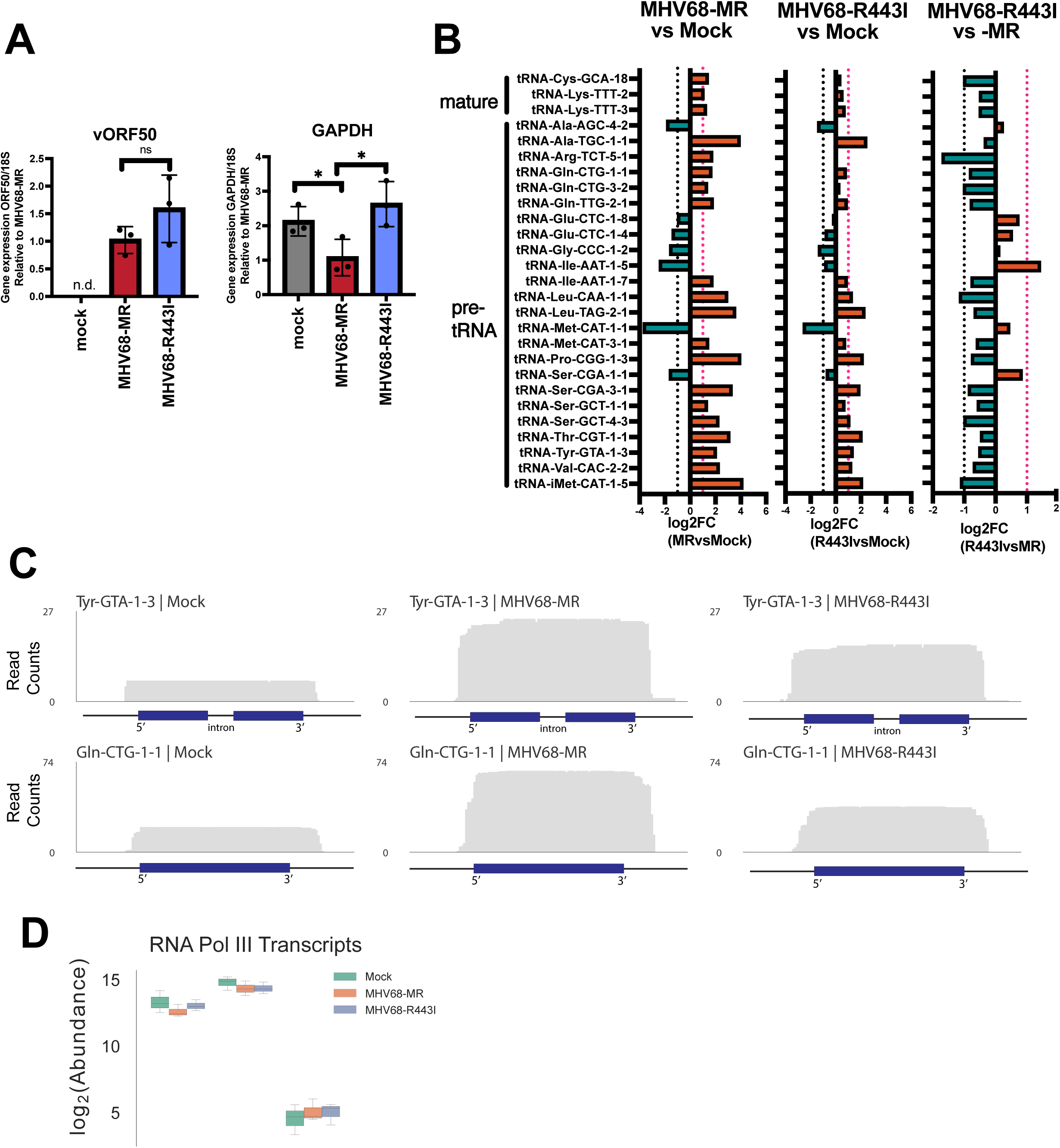
Pre-tRNA accumulation is dependent on MHV68-induced host shut off. (A) RT-qPCR was performed using total RNA from mock-, MHV68-MR, or MHV68-R443I-infected MC57Gs to detect viral ORF50 (vORF50) and host GAPDH transcripts. Data depicts means +/− SD from three independent experiments relative to the MHV68-MR sample, with p-values calculated using raw 1′Ct values and unpaired t-test. (B) Log2(fold-change) values of MHV68-MR infected vs mock (left) for significantly differentially expressed host mature and premature tRNAs (p<0.05) are graphed alongside the corresponding log2FC for MHV68-R443I vs mock (middle) and MHV68-R443I vs MHV68-MR (right). Horizontal dotted lines indicate a fold change of > or < 2. (C) Normalized read coverage (5’ -> 3’) from mock (left), MHV68-MR (middle), or MHV68-R443I (right) across the Tyr-GTA-1-3 (top) and Gln-CTG-1-1 (bottom) tRNA gene loci. Blue boxes correspond to the gene body of the tRNA, while the horizontal black lines correspond to the 30 bp upstream, 30 bp downstream, and/or intronic regions. (D) Box plot showing the log2(abundance) of three RNA polymerase III transcripts in our sequenced size class (50-200 nt) colored by infection conditions.

### tRNA fragments are generated during MHV68 infection and are dependent on host shutoff

Our primary motivation for applying OTTR-tRAX was to examine global changes in tRF expression during infection, as this has not been explored during DNA virus infection. OTTR-tRAX analysis of the 15-50 nt size class confirmed the production of tRFs in MHV68-infected cells (Fig 4A-left, MHV68-MR vs. mock infection, p<0.05, Supp Figs 1-3). Based on tRAX parameters, tRFs are binned according to the best match source sequence, with priority given to mature tRNA reference sequences, which were curated for the processed mature tRNA sequence complete with 3’ CCA tails^21^. tRAX assigns these tRFs as “5’ tRFs” if they map within 5 nt of the annotated 5’ end of the corresponding mature tRNA, “3’ tRFs” if they map within 5 nt of the annotated 3’ end of the corresponding mature tRNA, or “other” if they do not meet these parameters. In contrast, reads that contain leader, trailer, or intronic sequence do not map to the mature tRNA reference, and instead map to genomic reference sequences and were binned as “pre-tRFs.” tRFs as a whole showed higher expression in MHV68-MR compared to MHV68-R443I (Fig 4A-B, Supp Fig 1-3), with a few exceptions (see Glu-CTC-5 in Supp Fig 1). Overall, the dependence on muSOX activity was more striking for tRFs than observed for pre-tRNAs, as assessed by the difference in log2FC values presented in Fig 3B vs. Fig 4A. For many tRNA genes, there were distinct 5’ and 3’ pre-tRFs produced from the same transcript (Fig 4C). For example, pre-tRFs from the intron-containing tRNA-Tyr-GTA-1-3 and tRNA-Arg-TCT-5-1 genes originated from both the 5’ and 3’ exons, with extensions into nearby leader, intron, or trailer sequence. Both 5’ and 3’ pre-tRFs from these two tRNAs are expressed to a higher extent during MHV68-MR infection compared to either mock- or MHV68-R443I infection. Pre-tRFs were also observed from intron-less tRNA genes, such as tRNA-Gln-CTG (Fig 4C). However, pre-tRFs from Gln-CTG genes had more heterogenous ends compared to pre-tRFs from Tyr-GTA and Arg-TCT. Among the other tRFs upregulated in infected fibroblasts were 5’, 3’ and internally derived tRFs (Supp Figs 1-3). Almost all of the significantly upregulated 5’ tRFs had distinct 3’ ends, which resided in or around the D-loop (Supp Fig 1). In contrast, 3’ and internally derived tRFs exhibited more heterogenous ends, suggestive of ongoing exonucleolytic degradation (Supp Fig 2-3). In some cases, the tRFs could be mapped uniquely to a specific tRNA locus of origin (e.g., 3’ tRF-Asn-GTT-1 in Supp Fig 2 or int-tRF-His-GTG-1 in Supp Fig 3, purple reads). These results illustrate that MHV68 triggers tRF production, and that this phenomenon is dependent on host shutoff activity by MHV68 muSOX.

**Figure 4.**
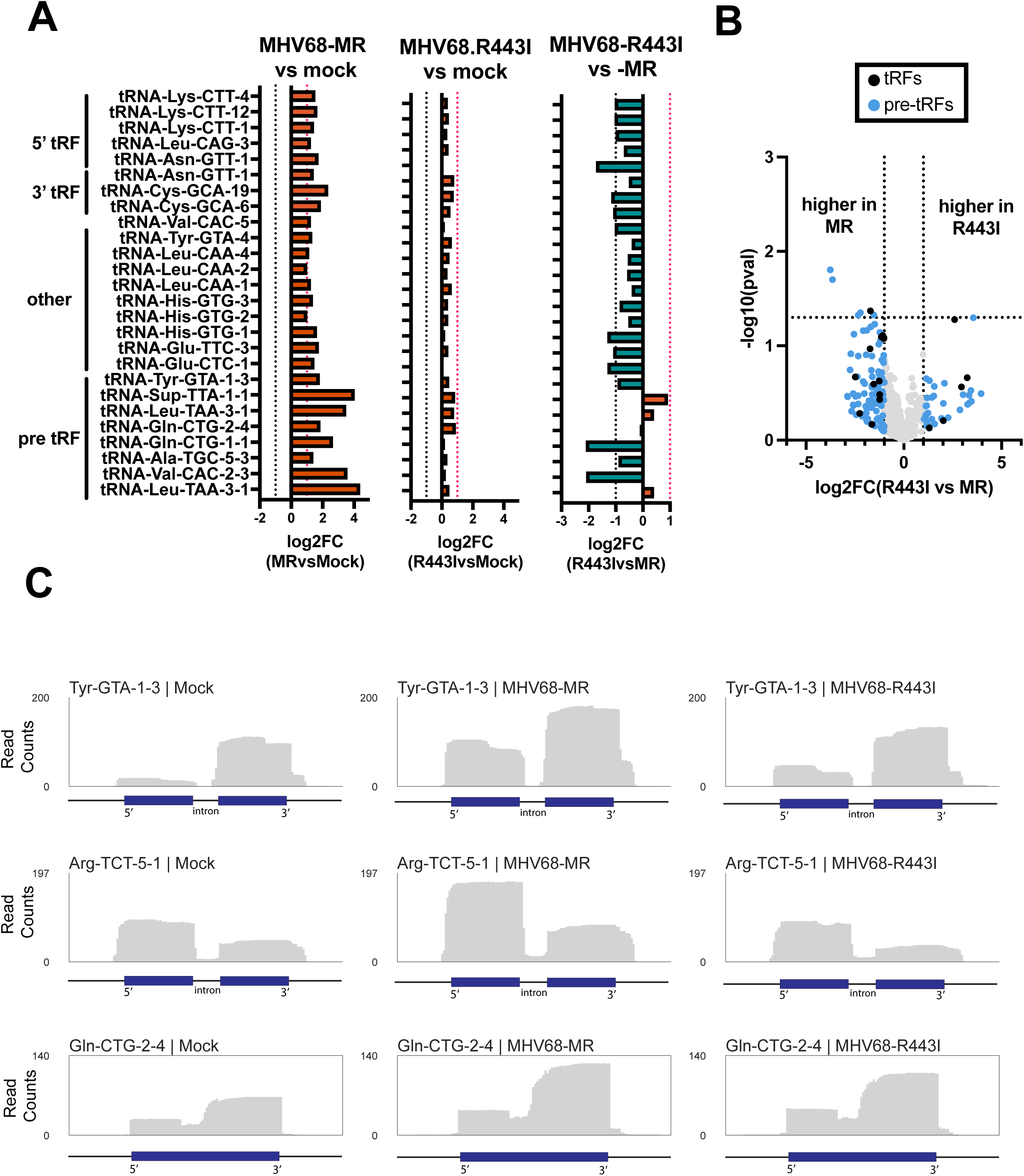
tRNA fragments (tRFs) are induced in a host shutoff-dependent manner. (A) Log2(fold-change) values of MHV68-MR vs mock (left) for significantly differentially expressed host cytosolic 5’tRFs, 3’tRFs, internal tRFs, and premature-derived tRFs (p<0.05) are graphed alongside the corresponding log2FC for MHV68-R443I vs mock (middle), and MHV68-R443I vs MHV68-MR (right). Horizontal dotted lines indicate a fold change of > or < 2. (B) Volcano plot of log2(FC) values for tRFs from MHV68-MR vs. MHV68-R443I infections, plotted against -log10(pval). tRFs up- or down-regulated more than 2-fold are colored blue for pre-tRFs or black for tRFs generated from mature tRNAs. (C) Normalized read coverage (5’ -> 3’) from mock (left), MHV68-MR (middle), or MHV68-R443I (right) across tRNA gene loci. Blue boxes correspond to the gene body of the tRNA, while the horizontal black lines correspond to the 30 bp upstream, 30 bp downstream, and/or intronic regions.

### Viral tRNAs are modified and undergo cleavage into tRNA fragments

We expected OTTR-tRAX might be a useful tool to examine changes in viral TMER and TMER-derived small RNAs during MHV68 infection. There are 8 encoded TMERs in the MHV68 genome, and TMER transcripts play an important role in pathogenesis and latency establishment during MHV68 infection in mice^13, 14^. Transcription of the TMERs by RNA polymerase III results in a hybrid RNA molecule consisting of a viral tRNA (virtRNA) fused to one or two miRNA stem-loops (Fig 5A). We use the “virtRNA” nomenclature here instead of the previously used “vtRNA” to avoid confusion with host vault RNAs. Processing of TMERs involves cleavage by the host enzyme ELAC2, normally responsible for removing the 3’ trailer during host tRNA maturation, to separate the viral miRNA stem loops from the virtRNA^15^. DICER then further processes the viral miRNAs for use by host silencing machinery. There is evidence that virtRNAs undergo maturation in the form of post-transcriptional -CCA addition at the 3’ ends^31^, which is a required step in host tRNA maturation to form the site of amino acid attachment. However, virtRNAs are not detectably charged with amino acids, at least for the virtRNAs previously tested (virtRNA3-6)^31^. We mapped OTTR-tRAX reads from both small and large-size classes to MHV68 TMER loci (Fig 5B,C). Our size selection at <200 nt excluded the majority of full-length TMER transcripts (∼200-250nt); however, we detected robust expression of virtRNAs from TMERs 1, 2, 4, 5, and 6 (Fig 5B). tRAX detects abundant misincorporations in transcripts, which reveal putative RNA modifications. We find 1-methyladenosine (m^1^a) modification-induced misincorporation at position 58A (seen as A>T in sequencing reads) in the majority of virtRNA transcripts detected (Supp Fig 4). Additionally, virtRNA5 contained near penetrant A>G substitutions, suggesting that virtRNA5 undergoes deamination at the wobble base position 34, resulting in inosine within the anticodon. To the best of our knowledge, this is the first demonstration that virtRNAs, similar to host tRNAs, can undergo modification and cleavage into stable fragments in MHV68-infected cells. For the smaller size class, we were able to detect the relative expression of viral miRNAs during infection in fibroblasts (Fig 5C). Interestingly, we detected fragments made from virtRNAs, with equal or higher abundance to viral miRNAs (see TMER4, 5, 7 as examples). TMER7 produced a highly abundant 3’ fragment, yet the virtRNA from this locus was not detected (compare Fig 5A and 5B). MHV68-R443I infection resulted in globally reduced expression of both virtRNAs and viral miRNAs compared to MHV68-MR, supporting previous microarray analysis^30^. Altogether, our analysis of TMERs revealed that virtRNAs undergo base modification and can produce shorter fragments, similar to host tRNAs.

**Figure 5.**
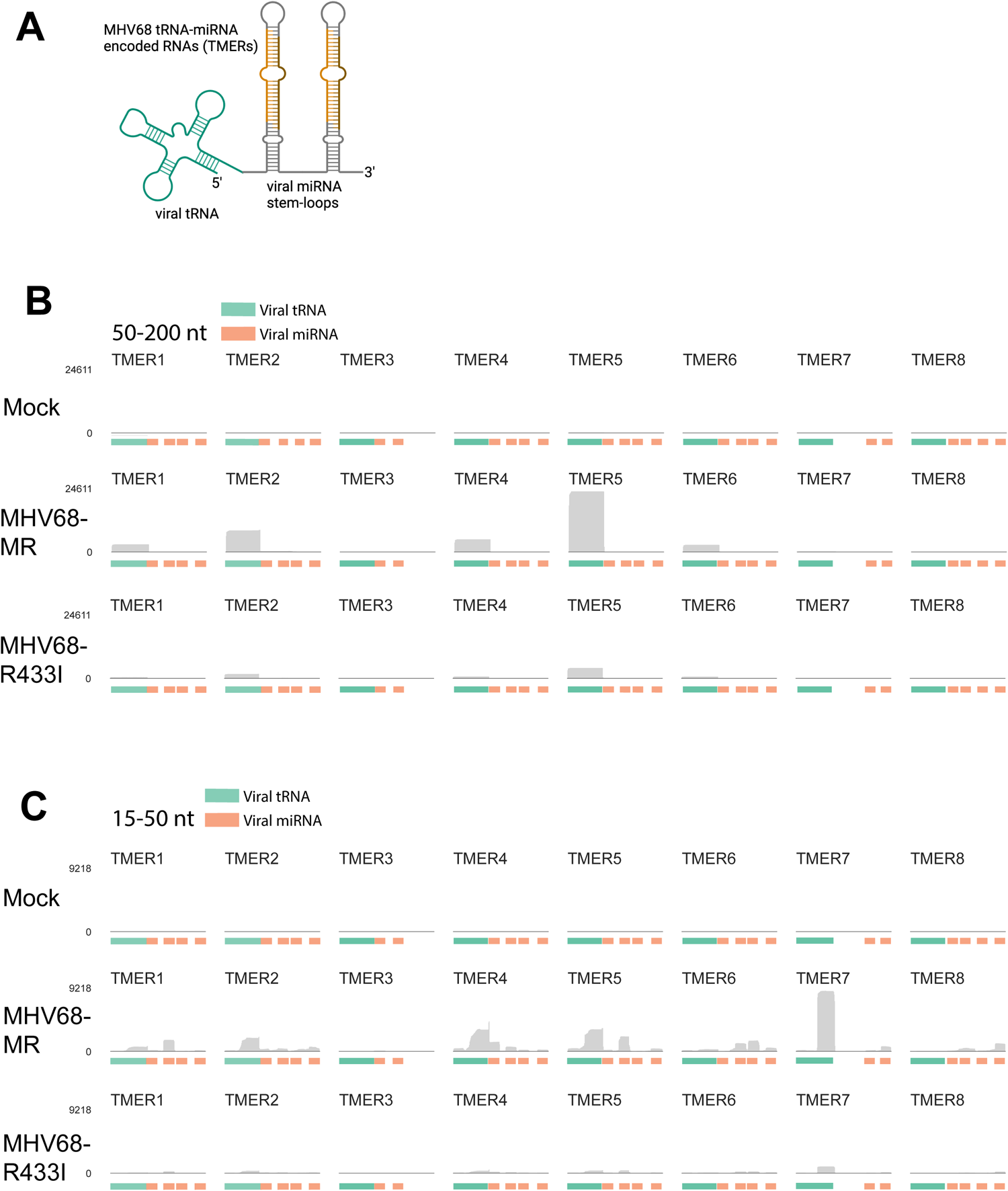
Viral TMERs are induced in a host-shutoff-dependent manner. (A) Schematic of a viral TMER highlighting the 5’ virtRNA and 3’ miRNA regions. (B) Normalized read coverage across the eight viral TMERs encoded within the MHV68 genome from the 50-200nt size selected libraries for mock (top), MHV68-MR (middle), and MHV68-R433I (bottom). (C) Normalized read coverage across TMERs from the 15-50nt size selected libraries for mock (top), MHV68-MR (middle), and MHV68-R433I (bottom). Color bars below indicate either a viral-tRNA feature (green) or a viral-miRNA feature (orange).

### tRNA fragments derived from pre-tRNAs are generated during MHV68 infection

We next explored potential biogenesis mechanisms of host tRFs. We noted that approximately one-third of tRFs produced in MHV68-MR-infected cells were derived from pre-tRNAs based on retained leader, trailer, or intronic sequences, leading us to hypothesize that upregulated pre-tRNAs are a major source of tRFs during infection. This was supported by our observation that MHV68-MR exhibited higher pre-tRNA accumulation and higher tRF expression, while MHV68-R443I showed lower pre-tRNA accumulation and lower tRF expression (Fig 4A-B). We reasoned that if tRFs are sourced from accumulating pre-tRNAs during infection, we would see a positive correlation between parental pre-tRNA and pre-tRF levels. We performed correlation analysis comparing log2FC(MRvsMock) for pre-tRFs vs. the parental pre-tRNA (Supp Fig 5) to assess the correlation between these two transcript classes. There was a weak positive correlation between isogenic pre-tRNAs and pre-tRFs, with a Pearson’s correlation coefficient (r) of 0.33 (pval=<0.001) and an r^2^ value of 0.11. This is not unexpected considering that steady state levels of pre-tRNAs and tRFs are dependent on stability features, including modification status and/or protein binding partners which might vary wildly between the two classes of RNAs. However, when we looked at each tRNA family individually, there were several tRNA families with strong, statistically significant positive correlation, including -Ala, -Asp, -Leu-, -Lys, and -Tyr (Supp Fig 5). Additionally, there was a strong positive correlation when we looked specifically at those tRNAs encoding introns (Supp Fig 5; r=0.77; p-val<0.0001; r^2^=0.60), which we posit is due to stronger mapping accuracy for pre-tRNAs and pre-tRFs. Together, we suspect that OTTR-tRAX underestimates pre-tRNA-derived tRFs, as internally derived tRFs lacking these identifying pre-tRNA features would be identical whether sourced from pre- or mature tRNAs, with tRAX prioritizing sequence mappings to mature tRNA reference sequences^21^. Regardless, there is a strong signature supporting that pre-tRNAs are substrates for endonucleolytic cleavage during MHV68 infection. Together, our data reveal a mechanistic link between pre-tRNA accumulation and tRF production.

### Validation of 5’ tRF production using stem-loop RT-qPCR

Next, we applied a method to detect 5’ pre-tRFs in response to MHV68 infection using an orthologous approach called stem-loop reverse transcription quantitative polymerase chain reaction (SL-qPCR). SL-qPCR allows the selective amplification of 5’ cleavage products versus their full-length counterparts^32^ (Fig 6A). Briefly, stem-loop reverse transcription (SL-RT) primers are designed to be specific for the 3’ end of the cleavage product. This SL-RT primer should preferentially bind to the cleavage product, not the precursor, as the precursor has a different 3’ end and additionally cannot be internally primed due to the steric hindrance with the SL-RT primer. We designed SL-RT primers for the upregulated 5’ pre-tRF-Tyr identified by OTTR-tRAX, as we had also validated its expression by northern blot (Fig 4A). We assessed the specificity of SL-qPCR for detecting 5’ pre-tRF-Tyr rather than full-length pre-tRNA-Tyr by using synthetic RNAs (Fig 6B). SL-qPCR was performed using a dilution series of equimolecular full-length and fragment synthetic RNAs as templates and confirmed >1000-fold specificity for fragment detection. We then applied the SL-qPCR assay for 5’ pre-tRF-Tyr to cellular RNA from cells +/− MHV68 infection (Fig 6C). We measured full-length pre-tRNA-Tyr using standard RT-qPCR and a reverse qPCR primer specific for the intron (absent in 5’ pre-tRF-Tyr) alongside as a control^6^. There was a 2.5-fold increase in 5’ pre-tRF-Tyr levels upon infection, which was equivalent to the increase observed by northern blot done in parallel (Fig 6C,D). We designed SL-qPCR assays for 5’ pre-tRF-Arg-TCT-3-1 and 5’ tRF-Gln-CTG-1/2 and confirmed their upregulation during MHV68 infection (Fig 6E). Thus, we validated 5’ pre-tRF expression revealed by OTTR-tRAX and established an effective molecular assay to monitor 5’ pre-tRF levels.

**Figure 6.**
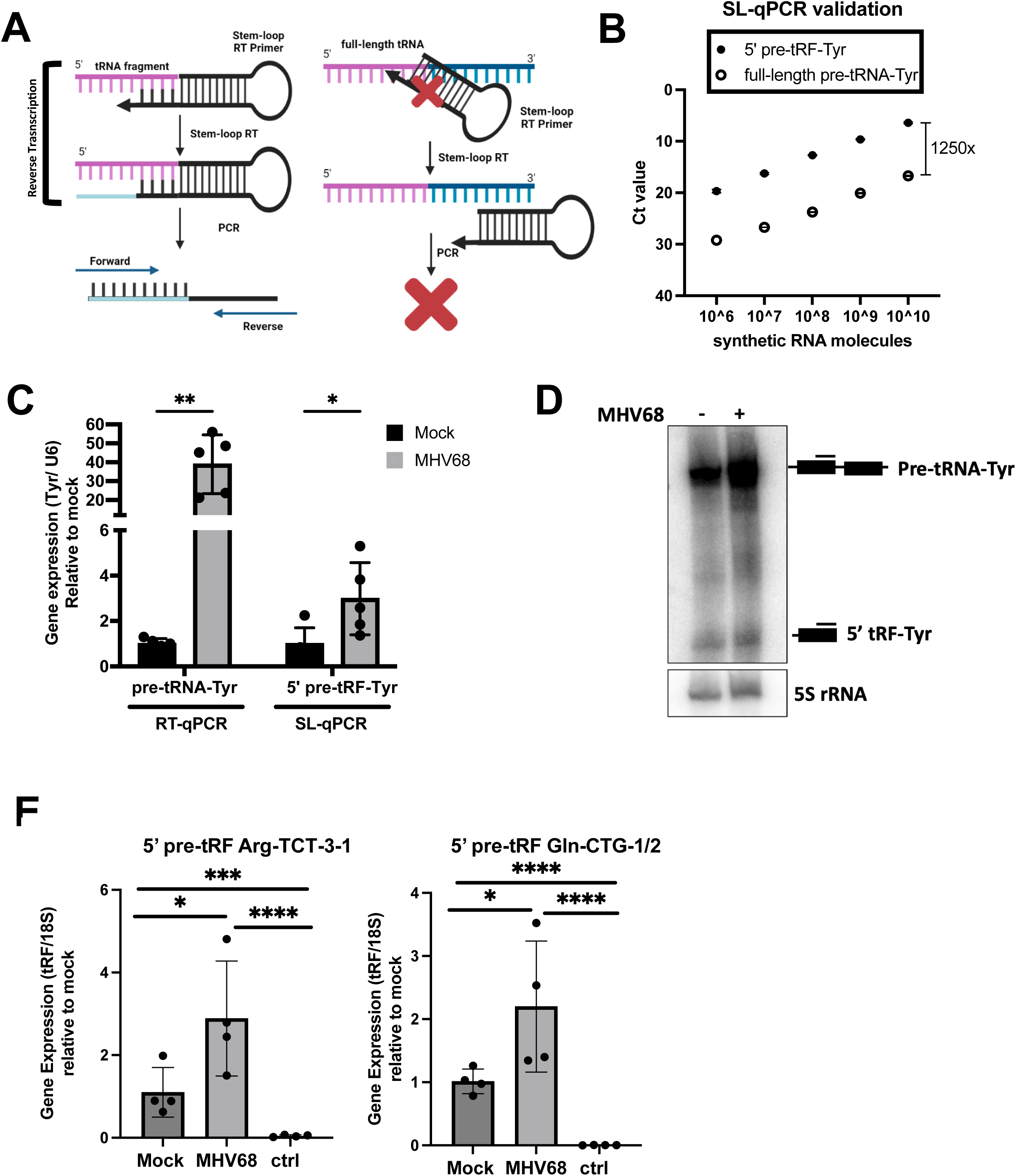
Stem-loop qPCR detection of 5’ tRFs. (A) Schematic illustrating the specificity of the stem-loop (SL) RT primer for 5’ pre-tRF vs. full-length parental tRNA. (B) Synthetic full-length pre-tRNA-Tyr or 5’ pre-tRF-Tyr were used in known quantities for SL-qPCR using Tyr forward and SL reverse primers. Raw Ct values are shown. (C) Standard- and SL-qPCR was performed using total RNA from mock or MHV68 infected NIH 3T3s to detect the full-length pre-tRNA-Tyr or 5’ pre-tRF-Tyr, respectively. Data depicts means +/− SD from three independent experiments relative to the mock sample, with p-values calculated using raw 1′Ct values and paired t-test. (D) Northern blot using 5’ exon Tyr probes was performed using RNA samples as described in (C). (E) SL-qPCR was performed from four independent experiments to detect 5’ tRF-Arg-TCT and -Gln-CTG as in (C). The sample labeled “ctrl” was reverse transcribed with a SL RT primer for an unrelated tRF species. tDRnamer^21^ nomenclature for amplified tRFs is reported in Materials and Methods. ns = p>0.05; * = p ≤ 0.05; ** = p ≤ 0.01; *** = p ≤ 0.001

### 5’ pre-tRF-Tyr expression is dependent on tRNA splicing factors

We noted that the 3’ end of 5’ pre-tRFs made from intron-containing tRNA genes (such as pre-tRNA-Tyr) aligned with the splice junction (Fig. 4C), suggesting that tRNA splicing may occur prior to or during pre-tRF biogenesis. Additionally, 5’ pre-tRF-Tyr upregulation was previously reported in fibroblasts and tissue extracted from kinase-dead *Clp1^K/K^* mice.^33^ CLP1 kinase associates with the tRNA splicing complex (TSEN2,15, 34, 54) and negatively regulates tRNA splicing^34–36^. We observed that the most abundant terminal ends of 5’ pre-tRF-Tyr (-TAGA, -TGTA, -GTAG) were downstream of the anticodon (GTA) and bordered the terminal nucleotide of the 5’ tRNA-Tyr exon (Fig 7A). The 3’ end -TAGA was the most abundant transcript termini observed in mock-infected cells (46.2%) and composed a slightly higher percentage in MHV68-infected cells (57.4%). Though 5’ leader nucleotides are present on 5’ pre-tRF-Tyr, the terminal -TAGA most likely results from cleavage of the anticodon loop post-splicing, as the terminus spans to the first nucleotide of the 3’ exon. This might suggest that intron removal can occur prior to trimming of the 5’ leader, which has not yet been described in mammalian cells, or perhaps solely is a byproduct cleaved from an aberrant tRNA precursor. In contrast, the -TGTA and -GTAG termini may potentially represent splicing intermediates.

**Figure 7.**
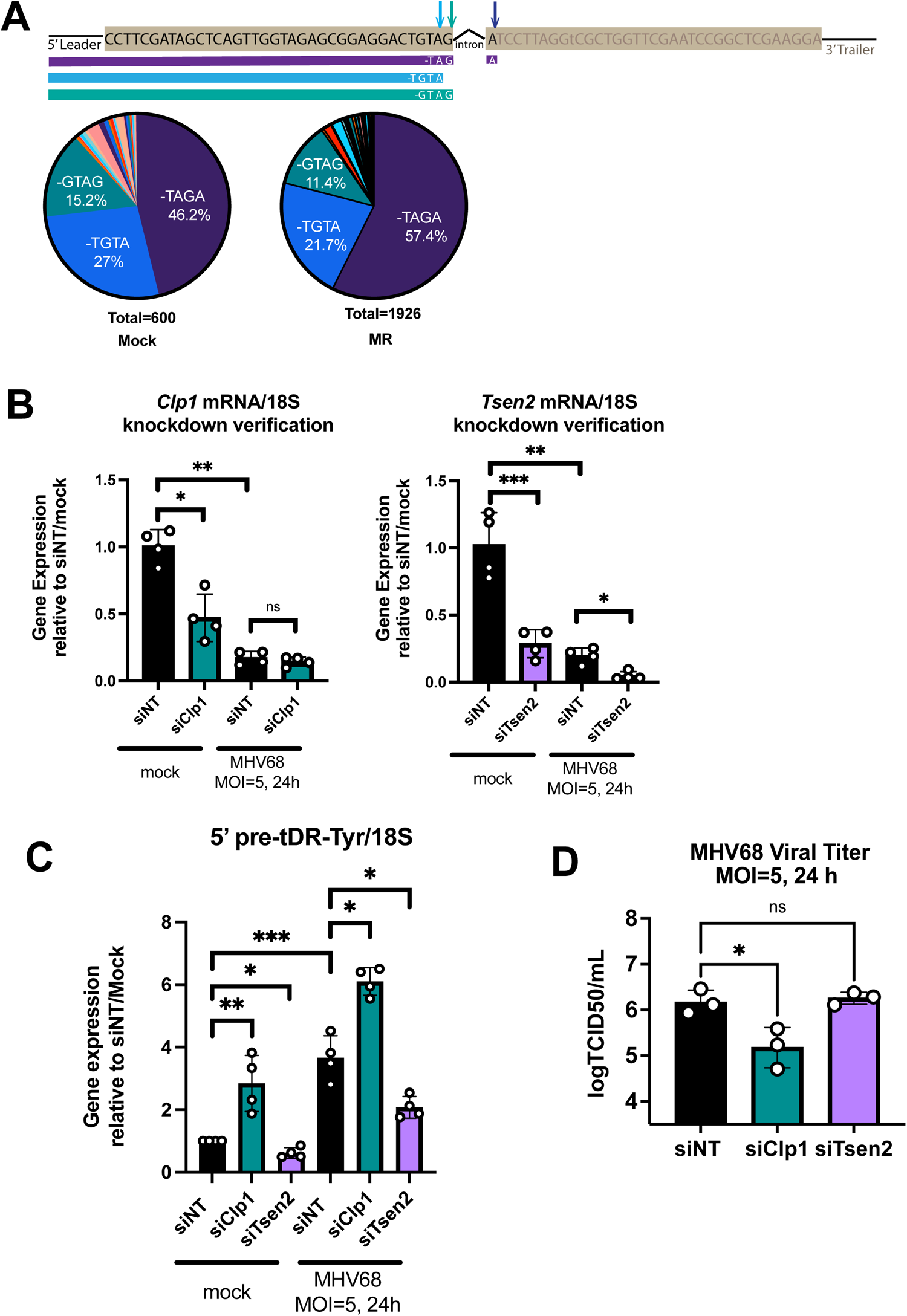
The tRNA splicing factors TSEN2 and CLP1 modulate 5’ pre-tRF-Tyr expression during MHV68 infection. (A) Schematic depicting a pre-tRNA-Tyr transcript (top line) and top three most abundant 5’ pre-tRF-Tyr transcripts detected (purple, blue, and teal lines). The pie charts depict the percentage of detected termini of pre-tRF-Tyr transcripts from mock- and MHV68-MR infected MC57G fibroblasts using OTTR-tRAX. (B) siRNA-mediated knockdown of *Tsen2* (siTsen2) and *Clp1* (siClp1) or treatment with non-targeting siRNAs (siNT) was followed by mock or MHV68 infection at an MOI=5 for 24 h. siRNA knockdown was confirmed using RT-qPCR using *Clp1*, *Tsen2*, and *18S*-specific primers. (C) SL-qPCR was used to measure 5’ pre-tRF-Tyr upon siRNA treatment. Data from RT-qPCR and SL-qPCR experiments is depicted as mean +/− SD from four independent experiments relative to the siNT/mock sample, with p-values calculated using raw 1′Ct values and paired t-test. (D) Viral titer of supernatants was measured by TCID50 from three independent experiments, with p-values calculated by one-way ANOVA. ns = p>0.05; * = p ≤ 0.05; ** = p ≤ 0.01; *** = p ≤ 0.001

The tRNA splicing endonuclease, TSEN2, was previously shown to be involved in 5’ pre-tRNA-Tyr production in *Clp1^K/K^* mouse embryonic fibroblasts^33^, and so we tested whether the same was true during MHV68 infection. To test whether 5’ pre-tRF-Tyr was sensitive to decreased expression of tRNA splicing factors, we used siRNAs to knockdown expression of both *Clp1* and *Tsen2* (Fig 7B). We validated knockdown of *Clp1* and *Tsen2* mRNA expression by RT-qPCR, as we were unable to obtain commercial antibodies of sufficient specificity (Fig 7B). Both knockdowns reduced expression of their target mRNA in mock-infected cells. Knockdown did not reach statistical significance for *Clp1* in MHV68-infected cells; however, *Clp1* and *Tsen2* mRNA expression was diminished due to host shut-off during infection and approached the limit of detection. *Clp1* knockdown was associated with enhanced 5’ pre-tRF-Tyr expression in both mock- and MHV68-infected cells, confirming that CLP1 negatively regulates the production of 5’ pre-tRF-Tyr in both contexts (Fig 7C). In contrast, *Tsen2* knockdown decreased 5’ pre-tRF-Tyr levels, suggesting that TSEN2 contributes to the accumulation of 5’ pre-tRF-Tyr in mock- and MHV68-infected cells, similar to *Clp1^K/K^* MEFs. Together, these data suggest that tRNA splicing may be dysregulated during MHV68 infection, leading to the production of pre-tRNA fragments. Finally, we measured infectious titer following a single round of MHV68 replication in cells treated with non-targeting, *Clp1*, or *Tsen2*-targeting siRNAs (Fig 7D). *Clp1* knockdown resulted in an 8-fold decrease in MHV68 titer compared to the non-targeting control, while *Tsen2* knockdown had no effect. These data suggest that CLP1 activity, perhaps through the control of tRNA splicing, supports MHV68 replication.

## Discussion

While there have been reports of tRFs produced in response to RNA virus infection^2, 4^, here we show that DNA viruses can also induce host tRNA cleavage. By combining OTTR-seq for library production with the bioinformatic software tRAX, we conducted the first in-depth analysis of tRNA and tRF expression during MHV68 infection. OTTR-tRAX offers a significant improvement over currently used tRNA sequencing strategies, as it: 1) utilizes a novel bioengineered RT that effectively copies modified tRNAs/tRFs and 2) minimizes sequencing artifacts. OTTR-tRAX facilitated the discovery that a variety of tRFs are upregulated in response to MHV68, with the majority derived from pre-tRNAs that accumulate during infection. Based on MHV68-MR and MHV68-R443I sequencing, we note a correlation between elevated tRF levels and the parental transcript of origin for numerous tRNA families. As viruses belonging to each of the three herpesvirus subfamilies (alpha, HSV-1; beta, HCMV; gamma, MHV68) have now been reported to induce tRNA transcription^6, 37, 38^, it will be interesting to examine if pre-tRNA cleavage is also a universal response to herpesvirus infection.

Our data suggest that the accumulation of pre-tRNA transcripts during MHV68 infection drives the production of tRFs. Approximately one-third of the tRFs produced retain sequences that confirm their derivation from premature tRNA substrates. Further, there is a general positive correlation between the differential expression of isogenic pre-tRNAs and their tRF products induced by MHV68. And most strikingly, we saw that decreased pre-tRNA accumulation with MHV68-R443I versus MHV68-MR resulted in decreased tRF expression. There is also reason to assume that OTTR-tRAX may underestimate the total number of pre-tRFs, given that pre-tRFs that do not retain leaders, trailers, or introns are indistinguishable from tRFs derived from mature tRNAs. Overall, it is possible that infection-induced tRFs binned as mature tRFs could in fact be derived from pre-tRNAs, and this is currently being investigated. It is possible that other tRNA mapping software and strategies that do not carefully bin pre- vs. mature-sequencing reads might also report tRFs that are made from pre-tRNA substrates. There are known examples of pre-tRF formation and functionality, including a 3’ trailer-derived tRF that sequesters La/SSB to block RNA virus infection^2^. Based on the abundance of tRFs made from pre-tRNAs we identified during MHV68 infection, it is intriguing to consider the possibility that tRFs produced during other infection or stress scenarios are also derived from pre-tRNAs, and that their functionality may be driven by pre-tRNA-specific sequence and hypomodification characteristics.

Viruses often encode “mimics” of host machinery to facilitate infection, and TMERs encoded by MHV68 are an intriguing example of this viral strategy. TMERs are transcribed by host RNA polymerase III and undergo maturation by host tRNA and miRNA processing factors, including Elac2 and Dicer^15^. Early studies revealed that the virtRNAs are not charged^31^, and thus virtRNAs have been considered as solely promoter sequence for the expression of downstream viral miRNAs^13^ or other recently discovered ncRNAs^39^ made from TMER loci. Our data emphasize how widely the expression levels of different virtRNAs vary, with virtRNA5 being the most abundant in fibroblasts. Surprisingly, 3’ tRFs are made from TMER-derived virtRNAs as well, with abundance greater than their downstream miRNA counterparts (e.g., TMER7). Our finding that full-length virtRNA7 is not detected by RNA sequencing aligns with initial predictions based on sequence gazing that this virtRNA would not form a functional tRNA^31^. Additionally, we found evidence that virtRNAs undergo modification, including m^1^a modification at position 58 (virtRNAs1,2,4,5,6,7) and deamination at the wobble base position 34 (virtRNA5). The m^1^a modification at position 58 is a highly conserved modification among all branches of life, and is known to be important for proper RNA folding^40^, while modification of the wobble base is essential for viability due to its requirement in relaxed decoding^41^. We argue that the presence of base modifications and novel processing events within virtRNAs suggest additional functionality yet to be described. TMER maturation should be explored further to expand our understanding of the functional capacity of TMERs and other viral non-coding RNAs.

Finally, our data suggests that tRNA splicing contributes to host pre-tRF formation in mouse fibroblasts during MHV68 infection. We noticed infection-induced pre-tRFs derived from pre-tRNA-Tyr and -Arg with 3’ terminal ends that matched or bordered the terminal ends of the corresponding tRNA 5’ exon. Because of this observation, we explored the possibility that these tRFs form due to dysregulated tRNA splicing events, as reported in Clp1 kinase-dead MEFs^33^. TSEN2 cleaves pre-tRNAs at the 5’ exon-intron junction, matching the third most abundant transcript end for 5’ pre-tRF-Tyr detected. We found that TSEN2 activity contributes to 5’ pre-tRF-Tyr production in both mock- and MHV68-infected cells, suggesting that 5’ pre-tRF-Tyr may be a splicing intermediate. Our results contrast with the 5’ pre-tRF-Tyr described in response to oxidative stress in the human mammary epithelial cell line, MCF10A, that is not dependent on TSEN2^42^. We posit that there are likely other endonucleases, outside of tRNA splicing endonucleases, responsible for the full repertoire of tRFs sequenced under different stress or infection conditions. We also tested the role of CLP1, a negative regulator of tRNA splicing^36^, and saw that *Clp1* depletion increased the levels of detectable 5’ pre-tRF-Tyr, regardless of whether the cells were infected by MHV68. Because *Clp1* knockdown, and not *Tsen2* knockdown, limits MHV68 infection, it could be possible that fine-tuning tRNA splicing, rather than the biogenesis of 5’ pre-tRF-Tyr and similar products, might be most functionally relevant during MHV68 replication. However, in light of the experimental evidence that tRF production is a cellular response to infection^2, 17, 43, 44^, is conserved from bacteria to humans^45–47^, and can functionally impact gene expression^7, 8, 10, 48, 49^, we favor the hypothesis that virus-induced tRFs are not solely byproducts but functional drivers of infection. Overall, we present a comprehensive profile of tRNAs and tRFs induced by MHV68 and highlight how tRNA processing can be manipulated by and support virus infectivity.

## Data Availability/Sequence Data Resources

Sequencing data was deposited in GEO: GSE255627.

## Acknowledgements

ACM, MMB, ARJ, HEU, and JMT acquired the data, as supervised by KC, TML, and JMT. ACM and JMT analyzed the data and prepared the manuscript. All authors edited and approved the final manuscript. We thank Lucas Ferguson for his contributions to the development and optimization of OTTR, as well as all present and past members of the Tucker, Lowe, Glaunsinger, and Roller labs for their helpful suggestions, discussion, and careful reading of this manuscript. Data presented herein were obtained at the Genomics Division of the Iowa Institute of Human Genetics which is supported, in part, by the University of Iowa Carver College of Medicine and the Holden Comprehensive Cancer Center (National Cancer Institute of the National Institutes of Health under Award Number P30CA086862).

---

**Supplemental Figure 1.**
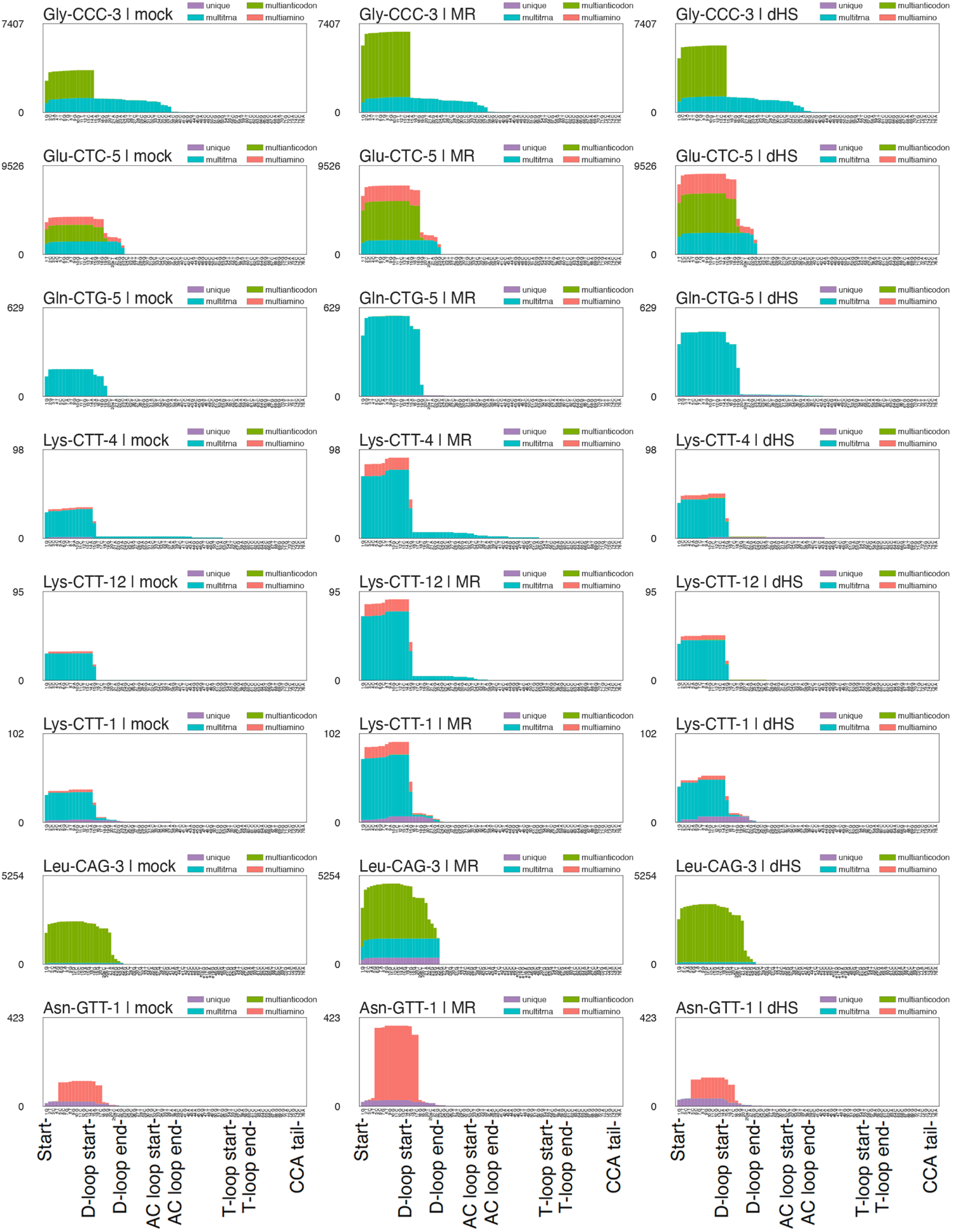
5’ tRFs induced by MHV68 infection. Normalized read coverage (5’ - 3’) from mock (left), MHV68-MR (middle), or MHV68-R443I (right) across tRNA genes. 5’ tRFs are defined by tRAX as reads that are within 10 base pairs of the start position, but do not reach the end of the full tRNA sequence. Colors of the coverage defines the specificity of read mapping. Purple “unique” reads uniquely map to the corresponding tRNA transcript sequence. Blue “multitRNA” reads map to the transcripts with the corresponding anticodon. Green “multianticodon” reads map only to transcripts of the corresponding tRNA isotype. Red “multiamino” are reads that map to more than one tRNA isotype.

**Supplemental Figure 2.**
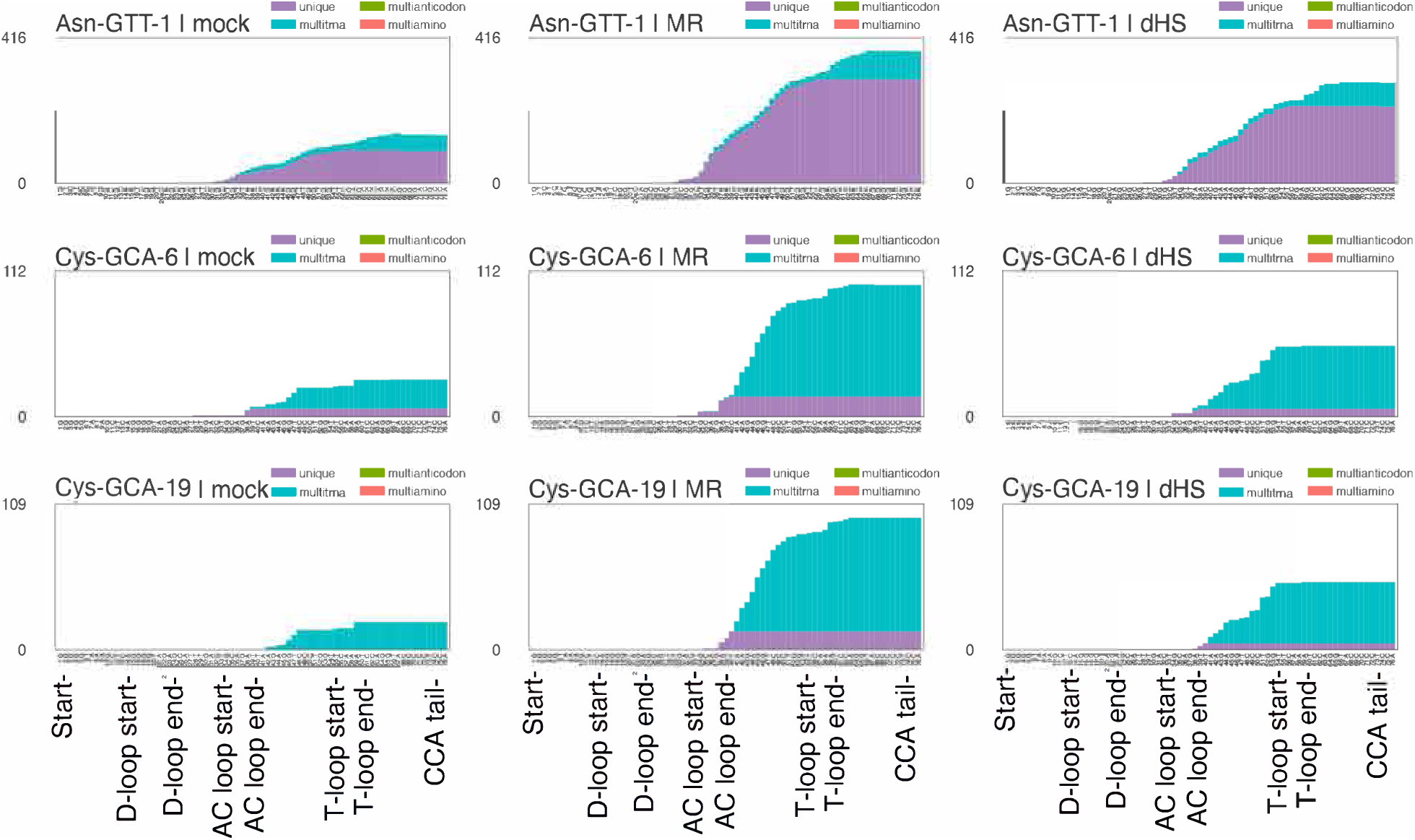
3’ tRFs induced by MHV68 infection. Normalized read coverage (5’ - 3’) from mock (left), MHV68-MR (middle), or MHV68-R443I (right) across tRNA genes. 3’ tRFs are defined by tRAX as reads that are within 10 base pairs of the end position, but do not reach the start position of the full tRNA sequence. See Supp Fig 1 legend for coloring information.

**Supplemental Figure 3.**
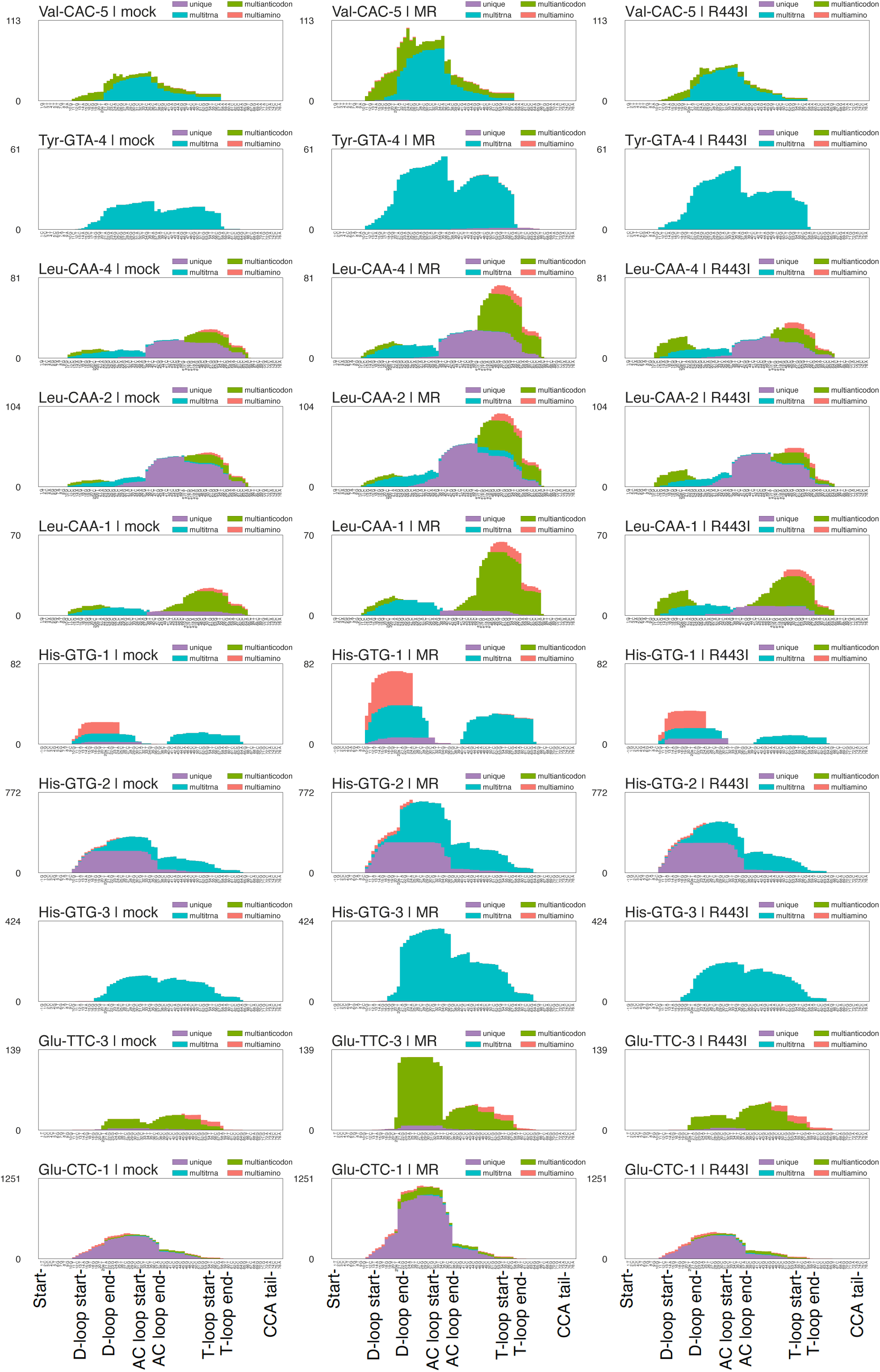
Other tRFs induced by MHV68 infection. Normalized read coverage (5’ -> 3’) from mock (left), MHV68-MR (middle), or MHV68-R443I (right) across tRNA genes. Other tRFs are defined by tRAX as reads that cannot be defined as 5’ or 3’ tRFs. See Supp Fig 1 legend for coloring information.

**Supplemental Figure 4.**
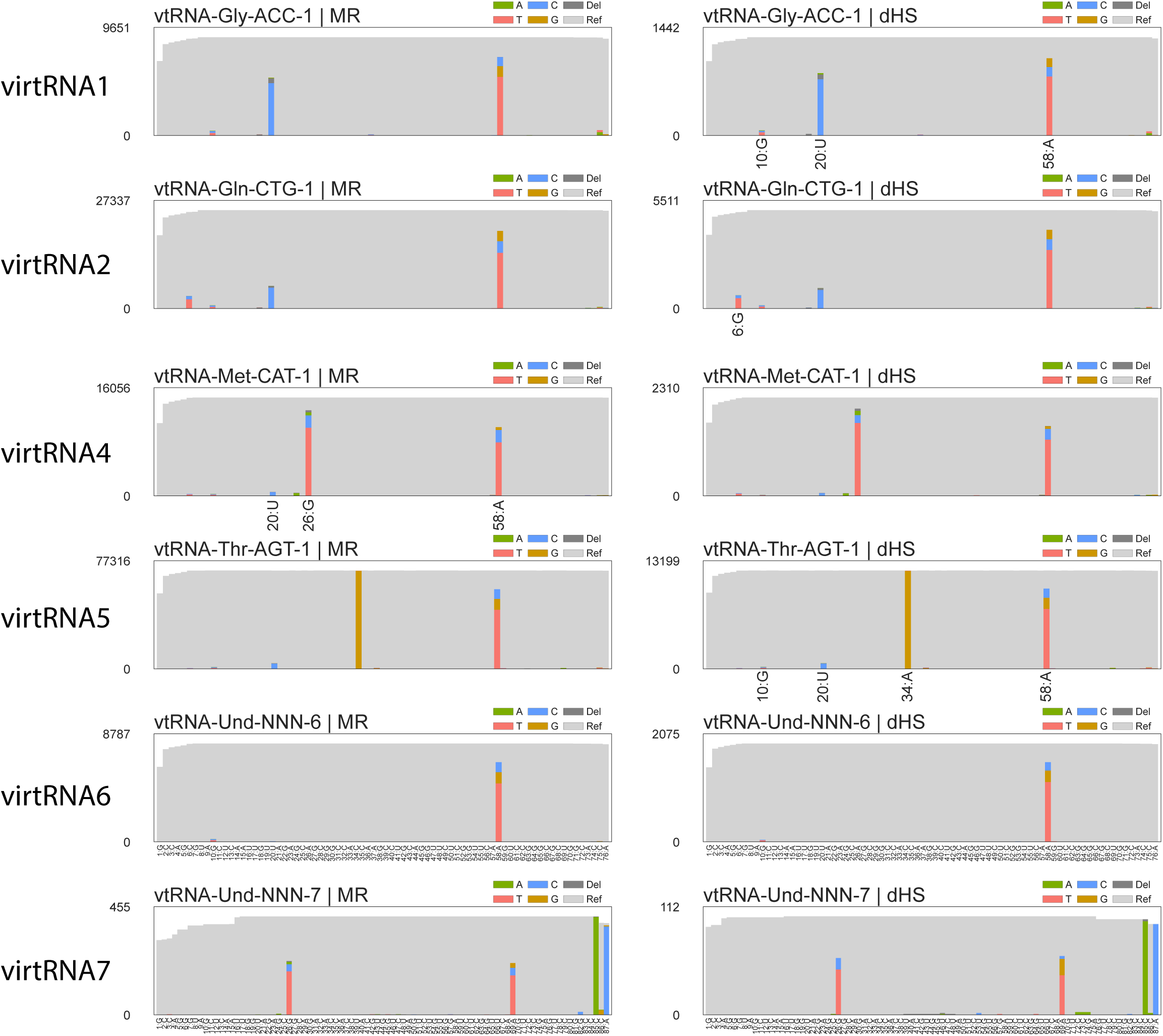
Viral tRNAs interact with host modification machinery. (A) Read coverage (5’->3’) across detectable virtRNAs. Grey indicates reads containing nucleotides matching the reference sequence, whereas further colors indicate the misincorporated base present. Sprinzl positions harboring a significant modification-induced misincorporation are highlighted.

**Supplemental Figure 5.**
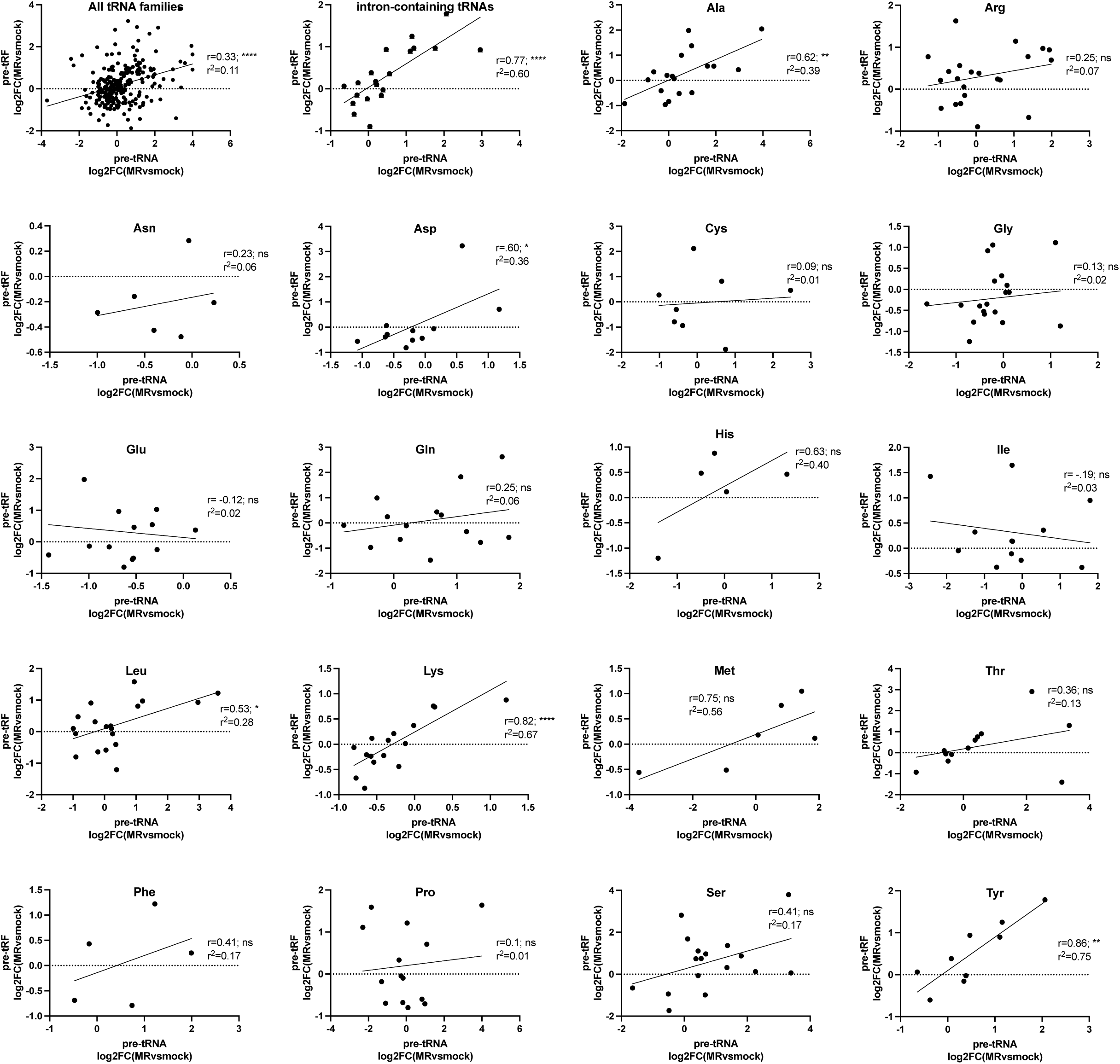
Correlative plots comparing pre-tRF expression to parental pre-tRNA transcripts. Log2(FC) values for pre-tRFs from MHV68-MR vs. mock infections are plotted against Log2(FC) values for their parental pre-tRNAs. The r^2^ value was calculated in Prism using the correlation function.

